# Recombination smooths the time-signal disrupted by latency in within-host HIV phylogenies

**DOI:** 10.1101/2022.02.22.481498

**Authors:** Lauren A. Castro, Thomas Leitner, Ethan Romero-Severson

## Abstract

Within-host HIV evolution involves latency and re-activation of integrated provirus that has the potential to disrupt the temporal signal induced by the evolutionary race between host immune responses and viral evolution. Yet, within-host HIV phylogenies tend to show clear, ladder-like trees structured by the time of sampling. Recombination complicates this dynamic by allowing latent HIV viruses to re-integrate as fragments in the genomes of contemporary virus populations. That is, recombination violates the fundamental assumption made by the phylogenetic methods typically used to study within-host HIV sequence data that evolutionary history can be represented by a single bifurcating tree. In this paper we develop a coalescent-based simulator of within-host HIV evolution that includes, latency, recombination, and population dynamics that allows us to study the relationship between the true, complex genealogy of within-host HIV, encoded as an Ancestral Recombination Graph (ARG), and the observed phylogenetic tree. We show how recombination recovers the disruption of the temporal signal of within-host HIV evolution caused by latency by mixing fragments of ancestral, latent genomes into the contemporary population through recombination. In effect, recombination averages over extant heterogeneity, whether it stems from mixed time-signals or population bottlenecks. Further, we establish that the signals of latency and recombination can be observed in phylogenetic trees despite being an incorrect representation of the true evolutionary history. Using an Approximate Bayesian Computation method, we develop a set of statistical probes to tune our simulation model to nine longitudinally-sampled within-host HIV phylogenies, finding evidence for recombination rates at the lower end of published estimates and relatively small latent pool sizes ranging from about 1000 to 2500 cells.

## Introduction

Within-host HIV evolution impacts between-host HIV evolution on the epidemic level [1, 2]. Thus, it is imperative to understand how within-host processes such as recombination and latency shape HIV phylogenies on different levels. Recombination violates the fundamental assumption that every offspring only has one parent (which is fundamental in a bifurcating tree). Instead, recombinant HIV taxa are formed by two parents in a generation, which cause loop structures in the resulting genealogical graph. Depending on the proportion and evolutionary history in each parent, this can lead to errors both in the branch lengths and implied evolutionary relationships among the sampled haplotypes [3]. Latency is the result of HIV provirus in dormant cells, which may be inactive for years, during which no evolution occurs. This may lead to extreme evolutionary rate differences [4], causing difficulties in tree reconstruction and especially in time-scaling trees. While these evolutionary effects are well-known in the HIV field, they are still often ignored simply because they are difficult to model.

On the epidemic level, the use of pathogen genetic data to infer epidemics has advanced considerably in the past decades. In such analyses the evolutionary history is nearly always thought of as a bifurcating tree, i.e., a standard phylogeny. Thus, such epidemiological applications [5, 6, 6–11] assume that 1) the genealogy of infection is a primary determinant of the phylogeny, and 2) the evolutionary history of a pathogen can be reasonably modeled as a bifurcating process, again, that the sequenced genetic regions are derived from one progenitor. Both of these assumptions are questionable. Within-host HIV evolution leads to substantial diversity, which has been shown to randomize transmission order and times [1, 12–14]. Thus, without modeling the underlying transmission history, the HIV phylogeny will be misleading about the details of transmissions. On a large population scale, where close donors and recipients unlikey are in the limited sample, however, the first assumption may be a reasonable approximation. The second assumption is less understood, and may thus present a bigger problem for the application of phylogenetic methods to HIV data.

Recombination is also fundamentally a problem for the epidemiological interpretation of HIV phylogenenies, although recombination at the epidemiological level sometimes is more obvious when it involves recombination of distinct so called subtypes with well known genetic signatures [15, 16]. The signal for teasing out recombination events is lower within subtypes, and even lower at the within-host level because many infections are established by a single virus [17, 18], which limits the extent of parental diversity that recombination detection methods use to identify recombinants. Likewise, the extent of recombination within-host leads to multiple generations of recombination events that further obscures the relationship between genealogy and phylogeny [19].

Estimates of the rate of HIV recombination within a host are very high. Both empirical [20] and simulation-based studies [21] estimate that the effective recombination rate, which incorporates both the probability of co-infection of a single host cell [22] and the template switching rate, is on the order of 1.4 × 10^−5^ to 1.38 × 10^−4^ per base per generation, comparable to the estimated point mutation rate of 2.2 − 5.4 × 10^−5^ per base per generation [23, 24]. Within a host, recombination provides HIV-1 with a means to increase the genetic variation for selection to operate on [25–29], can lead to rapid emergence of antiviral drug resistance [30–32], and can help shed deleterious mutations. Theoretical studies have shown that the magnitude of benefits brought by recombination depends on the interaction between factors such as population size and epistatic interactions [29, 33, 34].

Recombination clearly plays a central role in the evolutionary process of within-host HIV evolution, yet there are very few methods for actually modeling and addressing recombination empirically. Inference of more generalized recombination graphs called Ancestral Recombination Graphs (ARGs) has been proposed and demonstrated [35–37], but those methods are not scaleable to large tress [38]. Even if robust methods for inferring ARGs from data existed, there is very little understanding of how to interpret ARGs or how those results could be used to understand the interplay between recombination and other evolutionary processes. In this paper we develop a coalescent-based simulation method for simulating ARGs under a range of conditions including the rate of diversification, the level of recombination, the extent of latency, as well as within-host population bottlenecks in an idealized population intended to represent within-host HIV evolution. We implement a decomposition method to map ARGs simulated under a variety of conditions on to bifurcating trees. This method allows us to investigate the effects of within-host HIV biological processes on familiar phylogenetic trees without having to make unrealistic assumptions about the underlying genealogy. Combining the simulator with Approximate Bayesian Computation (ABC) methods and genetic data from nine serially sampled HIV patients, we also look at levels of recombination, latency, and population dynamics that are consistent with the real-world within-host HIV phylogeny.

## Methods

### Methodological overview

We simulate ARGs using a coalescent method that begins with a set of given samples at various times post-infection and simulates an ARG backwards in time according to a set of parameters intended to model the effects of recombination, latency, and selection. To map the simulated ARGs into the space of binary trees (i.e., what we can infer using standard phylogenetic methods) we decompose each ARG into a binary tree for each residue in simulated sequences by assuming a single random break point delineating the contribution from each parent. The average distance between tips is then computed over the population of decomposed trees and used to reconstruct a single tree that averages over the set of unobserved recombination events. Our ABC approach is based on comparing empirically measured statistics from real within-host HIV data to statistics computed on the simulated decomposed trees.

### Model Assumptions

We assume that an individual is infected with a single transmitted HIV-1 variant. From that lineage, the HIV-1 population diverges and diversifies linearly with time, with intermittent demographic bottlenecks caused by the host immune response. In addition, we assume that there is a reservoir latent population that is established three weeks after infection. The latent pool is assumed to be constant in size such that all movement to and from the latent pool is balanced. Integrated provirus in the latent pool does not acquire new mutations. For all processes, we assume neutrality, specifically that any lineage is equally likely to coalesce, recombine, go into or out of the latent reservoir, or survive a population bottleneck event. Lastly, we assume that the evolutionary rate remains constant over the course of the infection.

### Simulation of the ARG

We simulate ARGs in reverse time, beginning with the fixed set of sample times and moving backwards in time until the time of infection. All lineages have a state of being either *latent, L*, or *active, A*, and at any point in time there are *k*_*L*_ latent lineages and *k*_*A*_ active lineages. We model four possible events: (1) a recombination event between two active lineages, (2) a virus entering the latent reservoir, (3) a virus in the latent reservoir reactivating, and (4) a coalescent event between two active lineages. We assume that waiting times to each event are independent and conditional only on the extant number of lineages in either an active or latent state and the time since transmission. The model uses the parameters listed in Table 1.

**Table 1:**
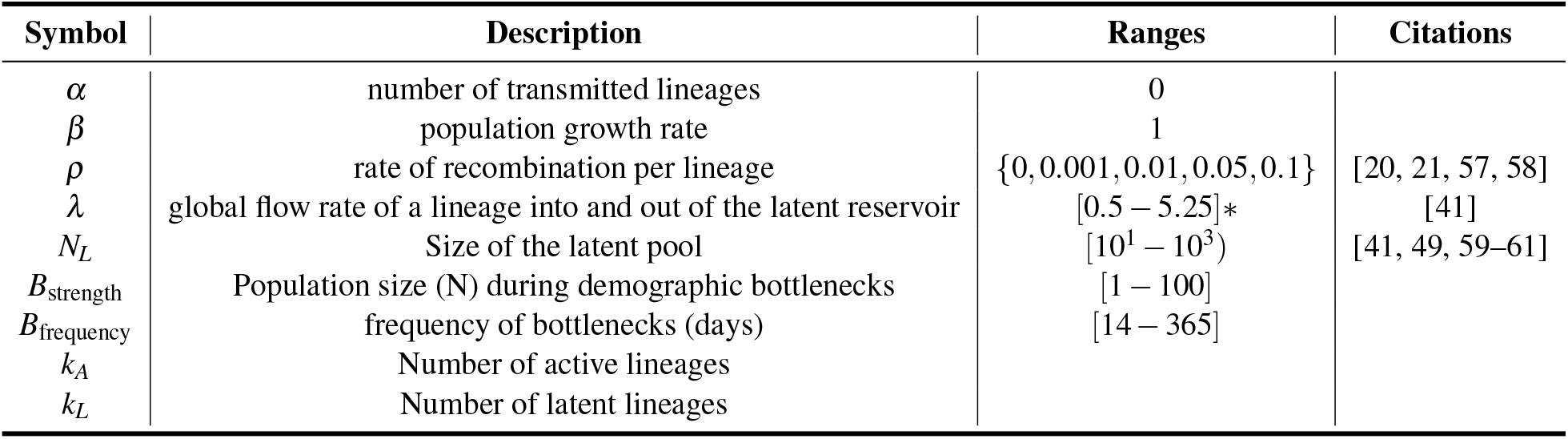
Parameter table. *This range of *λ* corresponds to these *N*_*L*_ values using the equation 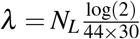.

### Simulation Events

- **Coalescence:** When a coalescent event occurs, two extant active lineages are chosen randomly and joined to form a single lineage (2*A* → *A*) and a coalescence node is inserted into the ARG. The expected waiting time for two random lineages to coalesce is dependent on the effective population size and the number of active lineages. The population size is modeled as N(t) = *α* + *β*t, where *α* is the number of transmitted lineages, *β* is the growth rate per generation (assumed to be 1 generation per day), and *t*, is the time since infection in days. For all simulations we assume that *α* = 0 and that *β* = 1. During bottleneck events, the population remains at a constant size, *B*_strength_, for a period of 5 days. Following the period of decreased population size, the population resumes linear growth, maintaining the overall gradual linear increase in genetic diversity over the course of the infection. In this way, we approximate the effects of selection without explicitly incorporating the reproduction probability of individual lineages. In the normal regime, the time to the next coalescent event is computed in the way described in [39]. During periods of bottlenecks that occur every *B*_frequency_ days, the time to the next coalescent event is an Exponential random variable with rate 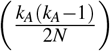[40].
- **Recombination:** A recombination event adds a lineage to the total number of extant active lineages (*A* → 2*A*). When a recombination event occurs, one lineage is chosen from the set of extant active lineages and is split into two lineages representing each parent. A node representing the recombination event is inserted into the ARG and a single breakpoint is assigned to that node with uniform probability over the set of simulated residues. We assume recombination to be a homogeneous process where recombination events occur at rate *ρ* per lineage, and thus the time to the next recombination event is an Exponential random variable with rate (*ρk*_*A*_).
- **Latent Reservoir Deposition and Reactivation**: The size of the latent pool, *N*_*L*_, is assumed to be constant with a fixed half-life of 44 months [41]. Therefore the per day rate of lineages moving into and out of the latent pool is 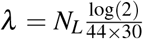; as the latent pool gets bigger, the rate into and out of the pool needs to increase to maintain a constant half-life. The overall rate of activation and deposition are equal but we also need to consider the probability that an activation/deposition event is ancestral to the sample. For activation (*L* → *A*), the time to the next event is an Exponential random variable with rate 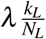. For deposition (*A* → *L*) the time to the next event is given by the same equation that governs the time to the next coalescent event with the 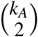 term replaced by *λk*_*A*_. Both deposition and re-activation events are recorded in the ARG as a node along a single branch indicating that the state of that lineage has switched.

#### Longitudinal Sampling

To simulate longitudinal sampling we designate times along the reverse time-axis when new active lineages are added to the simulation. At each additional sampling event, the branch lengths of any remaining active and latent lineages from the previous sample are extended in time up to the next sampling time. From there the simulation proceeds as before, with *k* and *k*_*A*_ updated to reflect the additional new samples. The simulation ends after the last sampling time (first in forward-time) when *k*_*A*_ = 1 before *t* = 0.

### Mapping the ARG to a single binary tree

To map the simulated ARG into what we might observe with standard phylogenetic methods we: 1) decompose the ARG into a population of binary trees, one for each residue in the simulated sequence (Fig 1), 2) compute an average distance matrix from the population of binary trees, and 3) compute a single phylogeny from the average distance matrix.

**Figure 1:**
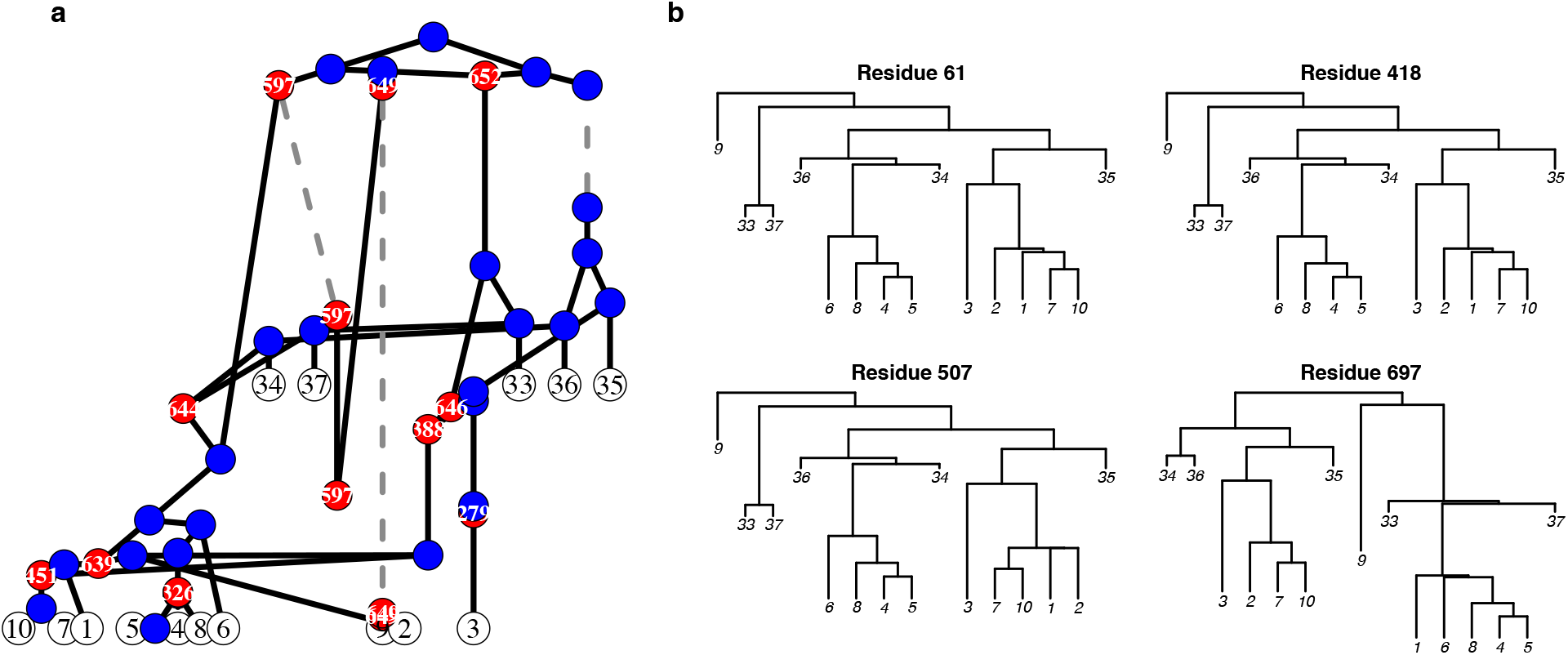
Ancestral Recombination Graph (ARG) decomposition into a series of bifurcating trees. (a) The simulated ARG from two sampling events. Red dots represent recombination events, with the break point identified in the white text. Blue dots represent coalescent events between two lineages. Dashed grey lines represent when a lineage is in the latent reservoir. Branch lengths correspond to time. This ARG represents the genealogical relationships among five viruses sampled 3.5 months post-infection and 10 viruses sampled 7 months post-infection and was simulated using *ρ* = 0.08, *N*_*L*_ = 760 assuming a sequence ength of 1000. (b) Each panel shows the decomposition of the ARG into a bifurcating tree for a specific position in the alignment. Recombinant (red) nodes are resolved at the nucleotide level by removing one of the parent branches depending on where the recombination event occurred with respect to the given alignment position. If no recombination occurred between two positions, then the genealogy for those positions will be identical (e.g. residues 61 and 418). However, recombination can alter both the topology and branch lengths of the underlying genealogy (e.g. residues 507 and 697).

For each residue, *i*, in the simulated sequence we extract a single binary tree by removing recombination nodes by trimming the right-hand lineage if *i* <= *y* where *y* is the breakpoint at that node and trimming the left-hand lineage otherwise. Because of computational limitations, we complete this decomposition process for a random sample of 25 residues.

For each binary tree we compute the distance between each tip by traversing the tree using the igraph package [42] in R and summing the total time along that path that was in the active state. When in the latent state, no evolutionary time is accumulated (i.e. we assume there is no potential for mutation in the proviral state).

Finally we use a minimum evolution principle [43] to generate a hierarchical clustering representation of the average distance matrix. To visualize a hierarchical clustered tree, we root it at MRCA of the samples from the first (in forward-time) sampling event. Hereafter, we refer to this as the *simulated tree*.

### Data

We applied this framework to the analysis of longitudinal HIV-1 DNA sequences sets from 9 patients [44]. In that study, sequences corresponding to the HIV-1 *env* gene were taken from each participant over a course of 6 to 12 years starting at 3 months from the time of seroconversion. There was an average of 11.875 individual sampling events with an average of 9.83 (std. 1.66) samples taken per event.

For our study, we aligned the sequences of each participant using MAFFT v7.305b2 and generated a tree using the balanced version of the FastME algorithm described in [43]. To translate the sequences into distance matrices we used the TN93 model [45]. The overall distribution of distances were similar using other evolutionary models. We obtained the sampling scheme from each participant, i.e. the number of samples (clones) taken at how many months post transmission. Thus for each participant, we have a distance matrix, a consensus tree, and a sampling scheme. To compare within-host evolutionary dynamics over the course of the same time duration, we restricted our analysis to samples taken no more than 90 months past the time of seroconversion.

### Selected statistics, objective function, and parameter inference

Our inference method is an Approximate Bayesian Sampling method using importance sampling to generate a population of parameters that produce simulated data that is most similar to the empirical data with respect to a set of statistics. To capture the general aspects of the empirical phylogneies we used 5 statistics (Table 2): the Sackin index (SI), the external to internal branch length ratio (EI ratio), the mean number of lineages though time (MLT), the coefficient of variation (CV), and the ranked pairwise difference (RPD). SI and EI are computed using standard methods. The MLT is calculated by the mean number of lineages in a clade with extant descendants across sample times. The CV is calculated by taking the sum of the absolute difference in the empirical and simulated standard deviation to mean ratio of pairwise distances across sample times. To map the simulations from time to genetic distance we assumed a static molecular clock optimizing the evolutionary rate to minimize the absolute difference between the simulated and empirical cophenetic distance matrices. The RPD is computed by taking the sum of the absolute difference in the empirical and simulated cophenetic distance matrices. To make the simulated and empirical distance matrices comparable, we ordered distances within each pair of sample times such that compared distances always have the same rank within a pair of time points. The objective function is then defined as the sum of each normalized statistic. If the objective function is 0 then the simulation has the exact same values of the measured statistics as the empirical tree for a given patient.

**Table 2:** Features used to match the simulated evolutionary history to a patient’s empirical HIV-1 *env* phylogeny. Some features measure tree characteristics (Topological) while others measure distance statistics (Distance). A simulation’s score, d, is the sum of the normalized differences between the feature measured on reconstructed simulated tree and the empirical tree.

| Feature | Feature Type | Description |
| --- | --- | --- |
| Sackin Index (SI) | Topological | The average number of splits from a tip to the root of a tree and captures asymmetry over the evolutionary history of the sample |
| Mean Lineages Through Time (MLT) [62] | Topological | The diversification of lineages normalized by the number of extant tips |
| External:Internal Branch Ratio (EI Ratio) | Distance | The ratio between the mean external branch length (branch that ends with a sampling event) and the mean internal branch length (branch between coalescence events) |
| Ranked Pairwise Distance (RPD) | Distance | The difference in the empirical and simulated distributions of branch lengths ranked by size within each sampling event |
| Coefficient of Variation in Pairwise Distance (CV) | Distance | The ratio between the standard deviation of pairwise distances between branches of the same sampling event and the mean pairwise distance between branches of the same sampling event |

To search for parameter sets that produced trees that are most similar to the empirical trees we drew 31,500 parameters sets within fixed strata of the recombination rate. Within each recombination strata parameters were sampled from [0.5 − 5.25] for *λ*, [1 − 100] for *B*_Strength_, and [14 − 365] for *B*_Frequency_, with uniform probability and assuming independence. We used the top 5% (i.e. with the lowest score) of parameters for each patient as the posterior distribution of parameters for each patient.

## Results

### HIV recombination and latency interact and affect the observed temporal signal in a within-host HIV phylogeny

To qualitatively investigate the interplay between recombination and latency, we first simulated a single ARG for a range of recombination rates and sizes of the latent pool of infected cells, and reconstructed the corresponding bifurcating phylogeny (Figure 2). Each tree has the same sampling scheme covering 132 samples over 147 months, which is representative of a very densely sampled within-host HIV phylogeny. The top left tree represents what happens when there is very little effect of latency–smaller latent pools have much lower re-activation and deposition rates to maintain a fixed half life of the latent pool–and no recombination. In this panel we see a tree that is orderly with respect to time (i.e., the tips sampled at the same time, more-or-less, occur at the same tree height). Moving along the top row (no recombination) left to right, the latency rate increases and we see almost immediately clear signs of temporal disordering: 1) large sets of sequences sampled at later times mixed with early samples, 2) lineages that appear to be extremely divergent from other sequences sampled at the same time, and 3) a collapse of the ladder-like backbone of the tree structure. Pattern 1 occurs when a period of latency happened on a branch that is ancestral to several tips that are sampled later, while pattern 2 occurs when little or no latency occurs on a single branch leading to an apparently highly divergent taxon. Pattern 3 is a result from randomly inserting ancestral variants from the latent pool, which destroys the basal time-structure. Thus, in the absence of recombination, even relatively modest levels of latency can completely disrupt the temporal signal in a phylogeny.

**Figure 2:**
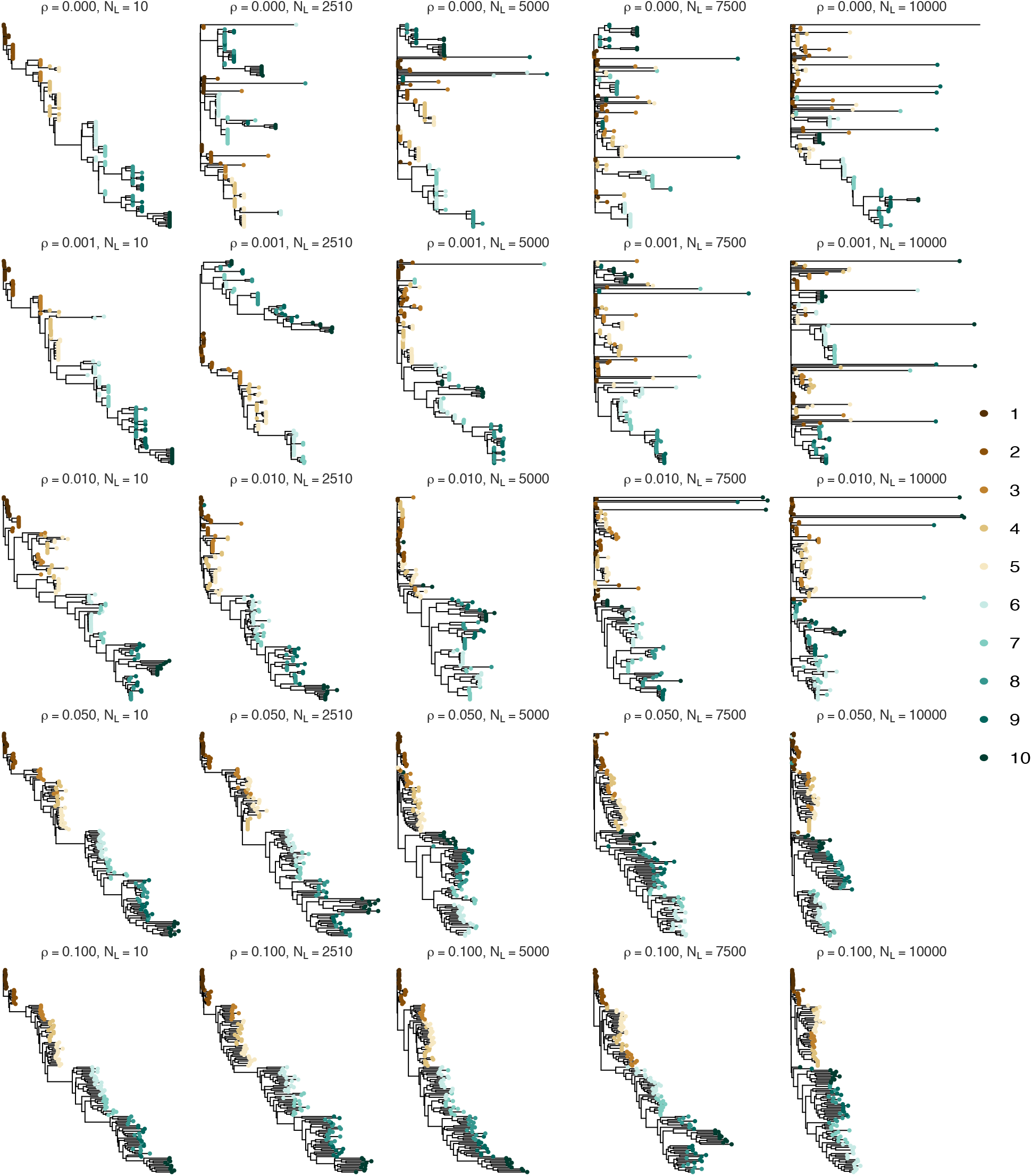
Simulated trees under a range of recombination rates and latent pool sizes. This figure shows a simulated within-host phylogeny under the specified recombination rate, *ρ*, and latent pool size, *N*_*L*_. Latent pool sizes increase left to right (a larger latent pool means more frequent deposition and activation of latent lineages) and the recombination rate increases top to bottom. The color of the tips represents the time from infection in years.

In the left column, moving from top-to-bottom, the recombination rate is increased and the level of latency reactivation is low. In general, even high levels of recombination don’t disrupt the temporal signal, however, it can cause some co-clustering of samples from different time points, but still overall has a distinct ladder-like structure. Interestingly, the higher levels of recombination in our study tended to produce longer external branch lengths, which could easily be confused as a signal for exponential growth. In general, moving from top to bottom, at all levels of latency, recombination clearly ‘recovers’ the temporal signal inherent in the data even at very high levels of recombination. At recombination rates above 0.05, we no longer see highly divergent lineages as we are only sampling recombinants of those lineages with other less diverged lineages (i.e. those with periods of latency in their past). However, when both the recombination and latency are high, it is still possible to see a slight disordering of the temporal signal as later sampled tips can co-cluster with earlier samples.

### Combined evolutionary effects generate the familiar HIV within-host phylogenetic structure

Our coalescent ARG simulator included several evolutionary processes that each could violate the fundamental assumptions on which standard phylogenies are based. As we have already seen in Figure 2, recombination rescues the disruptive effects of latency. This is surprising because recombination itself severely violates the bifurcation assumption. Thus, we next investigate this theme of recombination apparently amplifying, or at least rescuing, the phylogenetic signal disrupted by other evolutionary processes, also including periodic population bottlenecks.

Real within-host HIV phylogenies based on sequences sampled over time characteristically display unbalanced, ladder-like, time-ordered trees [44, 46]. With more frequent sampling over time lineages from different samples will overlap more, and conversely, over longer time only a few lineages survive from one sampling time to the next [47]. Simulating actual ARGs that include population demographics, recombination, and latency, makes it possible for us to investigate such effects on the expected bifurcating phylogeny that we are accustomed to see in HIV evolutionary research. We simulated within-host evolution over a 12 year period, sampled roughly yearly, to illustrate the effects that complex evolutionary processes have qualitatively on bifurcating trees. Overall, each of the evolutionary processes we simulate in our ARG framework show clear qualitative effects on the decomposed bifurcating trees (Figure 3).

**Figure 3:**
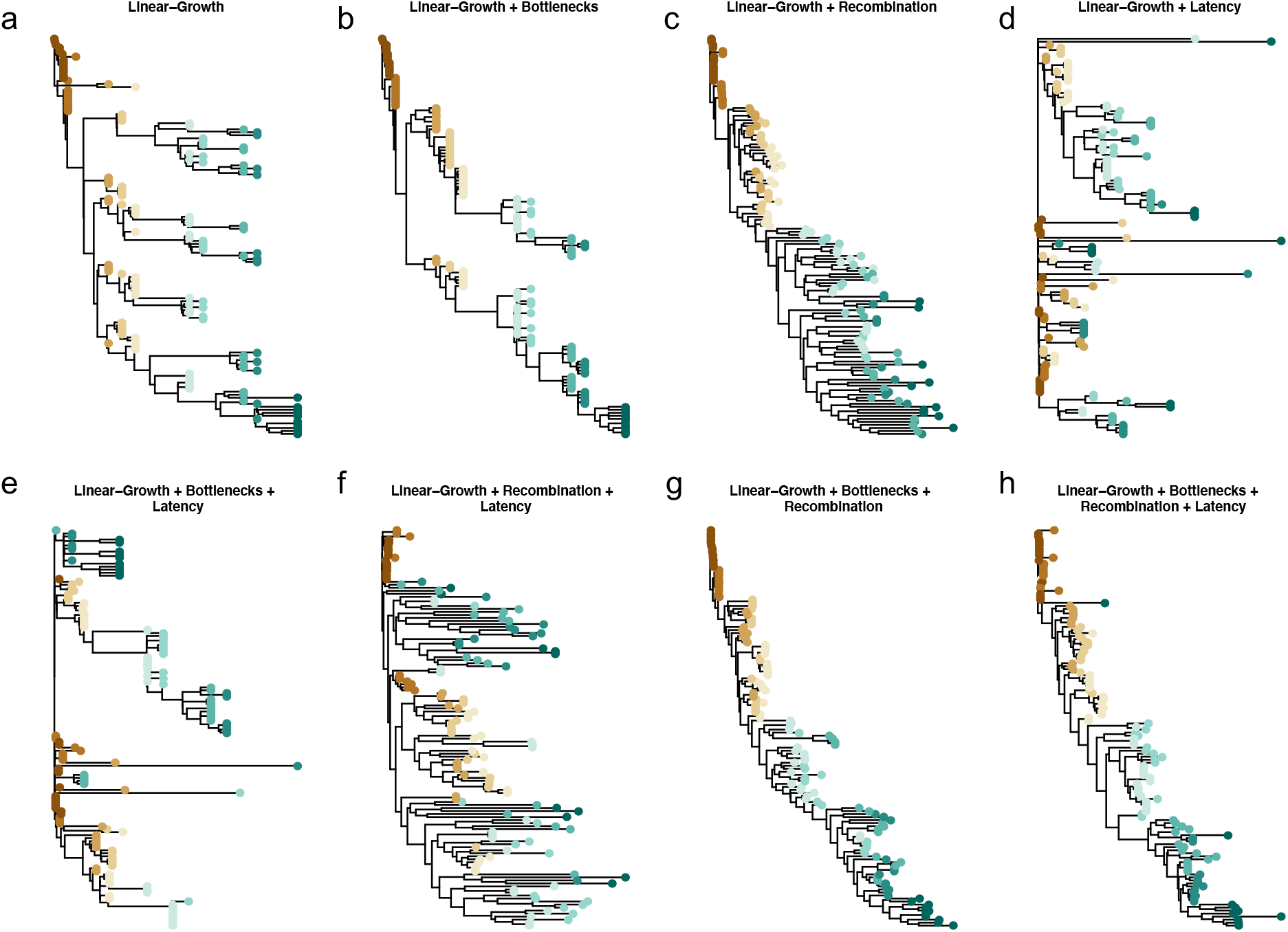
Illustration of effects of within-host evolutionary processes on virus phylogenies. Each tree is the decomposed average tree from one ARG simulation of 12 years using 132 sampled taxa spread over 10 sampling events. Color represents tips sampled at the same time. The baseline model, a linear-growth coalescent model is in panel A. Parameters are consistent across panels. For simulations in which the process is active: B_frequency_ = 300 and B_strength_ = 15; *N*_*L*_ = 2500; and *ρ* = 0.01.

In Figure 3a, we see an example of a tree generated using only a linear-growth coalescent model that accounts for linear increase in genetic diversity from the time of infection [39]. This model captures the lengthening of external branches as the infection progresses that is observed in empirical within-host HIV phylogenies. However, the long-term co-existence of distinct clades produced by simple linear-growth coalescent models are inconsistent with reality. While some HIV infected patients’ HIV populations can show multiple lineages surviving over time [48], this tree has too many parallel lineages co-existing and diversifying to appear fully realistic. As expected, adding bottlenecks to the simulation cuts down on the number of co-existing lineages (Figure 3b) as not all co-existing lineages will survive a bottleneck event. Surprisingly, adding only recombination to the linear growth produces trees that look plausibly like within-host HIV phylogenies (Figure 3c). This is due to the fact that recombination has the effect of averaging over extant heterogeneity (i.e. the co-existing lineages are intermixed causing them to co-cluster with one another), which produces trees that look like they have the signal of weak selection, but, importantly, do not. Without either population bottlenecks or recombination to trim and integrate the re-emergence of latent viruses (Figure 3d), phylogenies simulated with linear growth and latency alone do not resemble within-host HIV phylogenies at all. With latency in the model, adding bottlenecks alone cannot recover the expected tree shape (Figure 3e), while recombination can (Figure 3g). The tree in Figure 3f shows the characteristic co-clustering of heterochronous samples that is expected of a long period of evolutionary latency, but the branches appear very long suggesting long periods of population growth uninterrupted by selection. The final two trees (Figures 3g & h) show how these evolutionary effects, working simultaneously, can produce HIV phylogenies that very closely resemble real phylogenetic trees observed from longitudinally sampled patients.

### HIV recombination and latency have quantitative effects on phylogenetic tree features

Next, we investigated quantitative effects on the features used to match simulated and real HIV phylogenies (Figure 4). All of the statistics are variable under a broad range of both the recombination rate and latent reservoir size. That is, these features clearly respond to both of these important biological processes suggesting that, collectively, they form a reasonable probe for an ABC-based inference of these biological parameters. The empirical distributions of the feature statistics for 9 HIV-1-infected patients is shown in Figure 5. All of the selected statistics show high levels of heterogeneity between patients, possibly suggesting variability in both recombination rate and latent reservoir size between patients. However, Figure 4 also shows that stochastic effects produce fairly wide distributions in the feature statistics, which could also explain some of the between-patient heterogeneity in the within-host phylogenies. Interestingly, higher recombination rates reduce the variance in all the feature statistics, which is most pronounced in the external-internal branch length ratio. This is expected because recombination, in general, averages out differences between contemporary viruses at the sequence level, and therefore should reduce stochastic effects between simulation runs. Overall, both our method of simulating ARGs, decomposing them into bifurcating trees, reconstructing a consensus tree, and measuring the feature set of statistics on those trees is valid in that it produces self-consistent results that are linked to biological processes through our ARG simulator.

**Figure 4:**
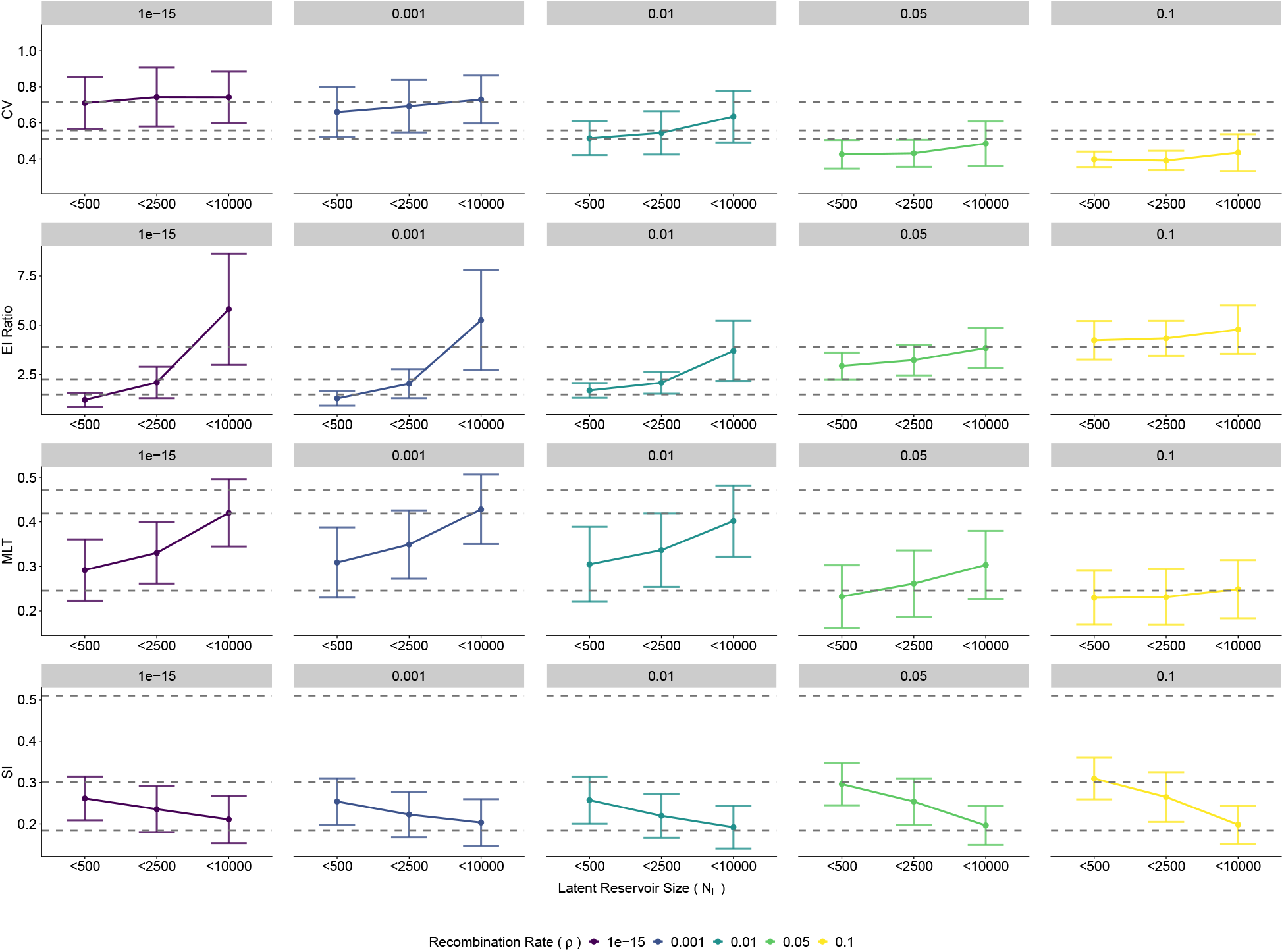
The magnitude of each biological process causes variability in distance and topological statistics. Each dot represents the mean and standard deviation of the statistic. Colors represent different rates of recombination, *ρ*. Simulations are grouped into three levels of reservoir size of increasing magnitude, with the breakpoints depicted on the x-axis. This figure shows all simulations where *B*_frequency_ < 200 days, and *B*_strength_ < 80. The grey dashed lines indicate the maximum, median, and minimum value for the 9 empirical patients in [44]. On the y-axis, the absolute limits of the MLT and the SI are from (0,1). We have shown that both the empirical and simulation values are a subset of this possible range.

**Figure 5:**
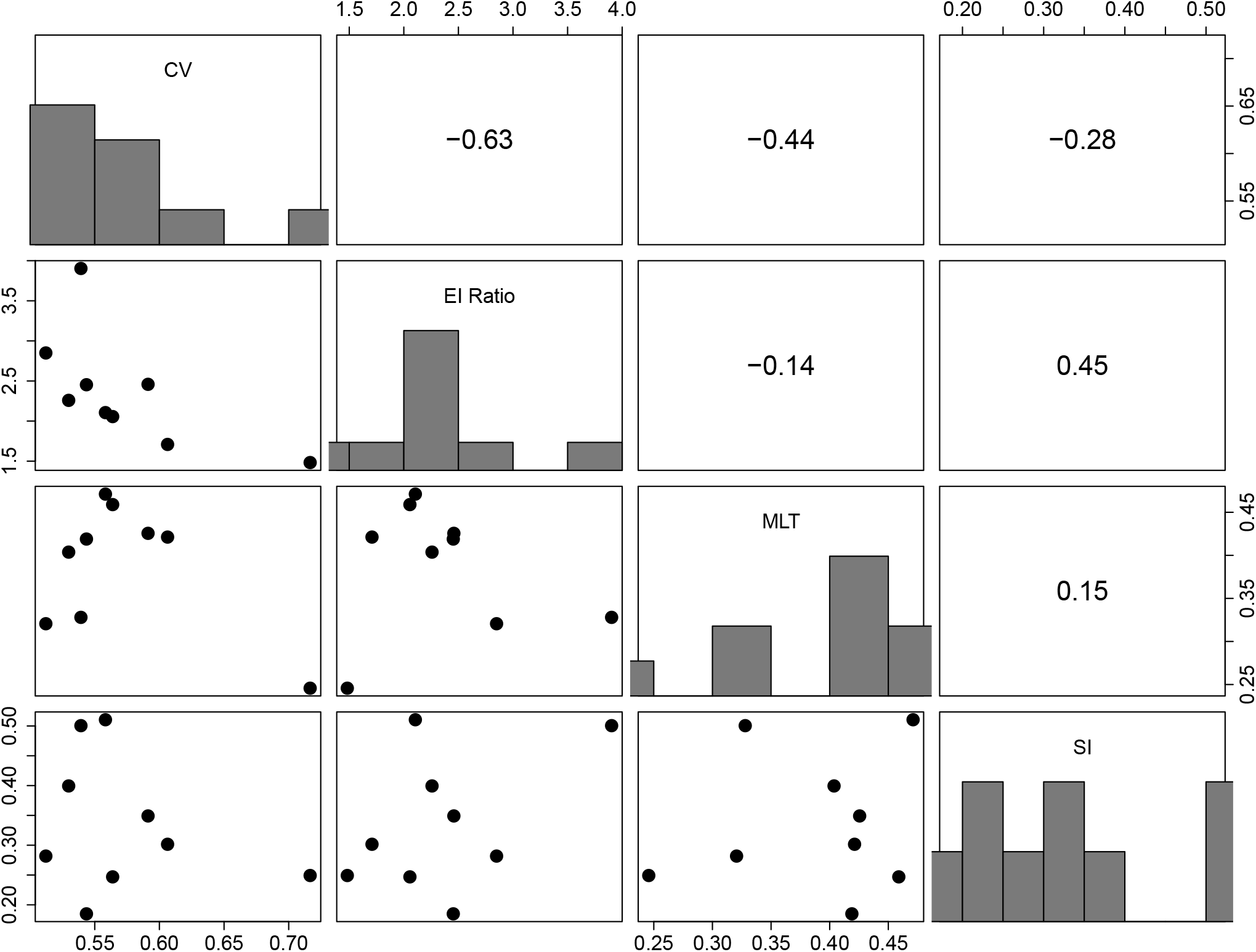
Feature values for the nine patients. These features were used to match simulations to the patient observed HIV-1 phylogenies. The CV captures the relative variability in the pairwise distance of tips from the same sampling event; the EI ratio is describes the relative differences between external and internal branches, the MLT calculates the mean number lineages of lineages in a clade with extant descendants across sample times, and the SI is an indicator of tree balance. We used an additional feature, the RPD; however we do not depict it here as it is a relational metric calculated directly between a simulated distance matrix and the observed empirical distance matrix.

### Application of ABC method to real HIV-1 within-host patient data

The simulator was also able to produce trees that resembled real patient data by minimizing the distance between the observed and simulated feature statistics. Figure 6 (panel B) shows two examples of observed and matched simulated trees. The simulated tree with the best score for patient 5 shows several of the typical features of the empirical tree including samples from specific time points at similar height and the general ladder-like tree structure with increasing external branch lengths in later time samples. While the simulator worked well for most of the patients, several of the patients showed overall low scores in the posterior samples (Figure 6, panel A) suggesting that their results are not as reliable (e.g. p9 and p11). The signal of recombination in the posterior samples for each patient is shown in Figure 7. For all patients, excluding p9, the recombination value of 0.01 recombination events per lineage per generation represented the plurality of recombination values in the posterior. Across patients, 0.01 represents an average of 52% of the posterior, and second most common recombination value of 0.001 represented on average 19.4% of the posterior. Overall, the majority of the posterior density is at recombination rates that are substantially lower than previously estimated values. The highest recombination rates reflecting typical published values were almost never in the top 5% of simulations suggesting that recombination rates above 0.05 are simply not compatible with observed phylogenetic trees. The size of the latent reservoir was generally well constrained by the data and showed only modest levels of heterogeneity between patients (Figure 8). Excluding patient 9, which appears to be an outlier in terms of the simulator, the mean latent reservoir sizes were between 1000 and 2500 cells, suggesting only a relatively small number of latency reactivation events were required to explain the empirical phylogenies. Interestingly there is a positive correlation in the recombination rate and the latent pool size (Figure S1), suggesting that if the latent pool is very large, then recombination must be rapidly integrating components of latent viruses into contemporary viruses. In general the bottleneck strength and frequency were not very strongly identified other than that extremely strong (i.e. perfect) and extremely frequent bottlenecks were ruled out for all patients (Figure S2), which is consistent with our observation that recombination alone even in the absence of bottleneck events can produce trees with the characteristic temporal ladder-like trees seen in within-host HIV phylogenies.

**Figure 6:**
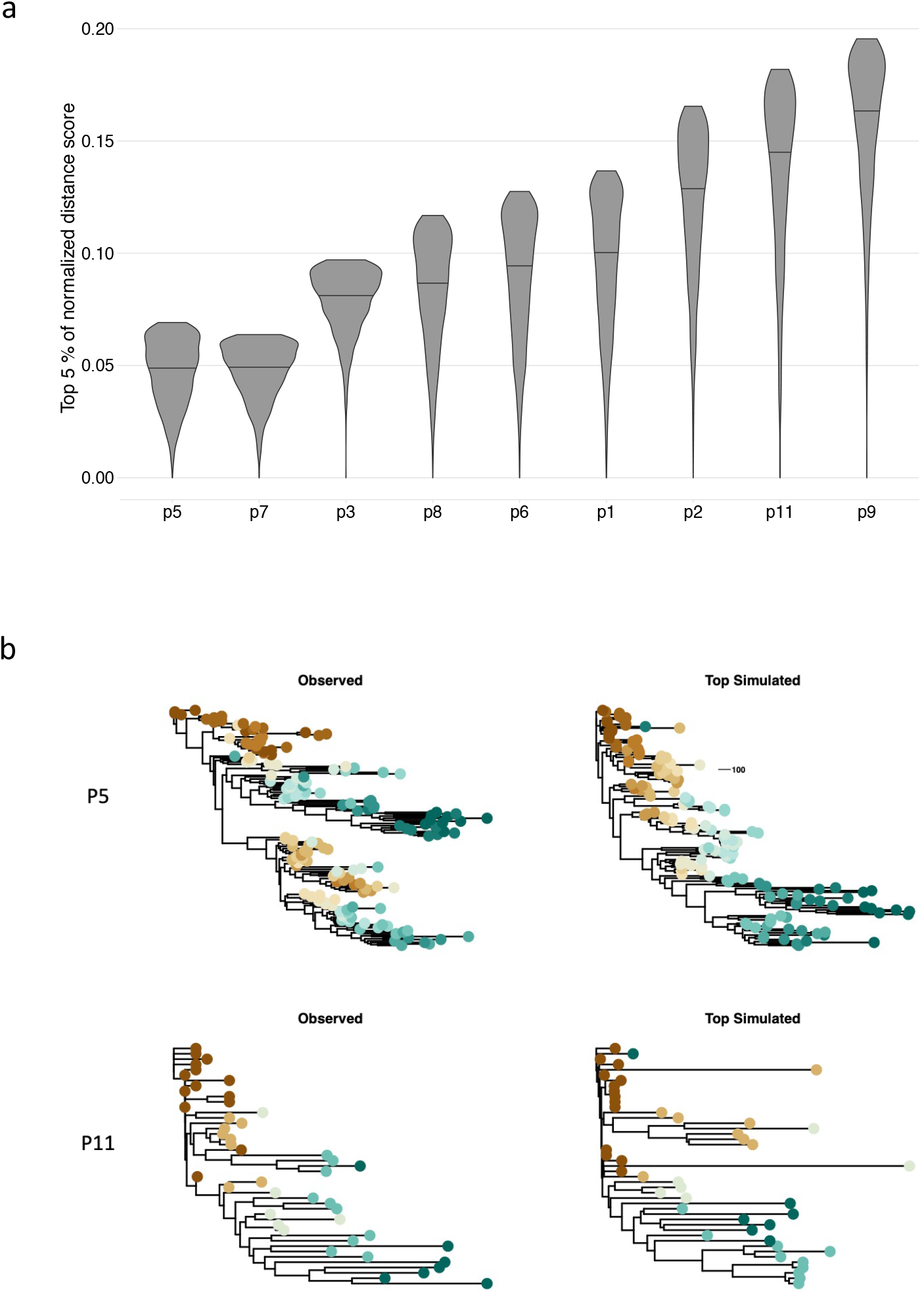
Normalized distance scores of the best-fitting 5% of simulations for each patient. Panel (a) shows the density of the best fitting 5% of 31,100 distance scores normalized for each patient, with the best-fitting match receiving a score of 0 and the worst-fitting match receiving a score of 1. Patients are ordered by the mean of the normalized top scores. Panel (b) illustrates the variation in quality of model fits by showing the observed and single best simulation for patients 5 and 11.

**Figure 7:**
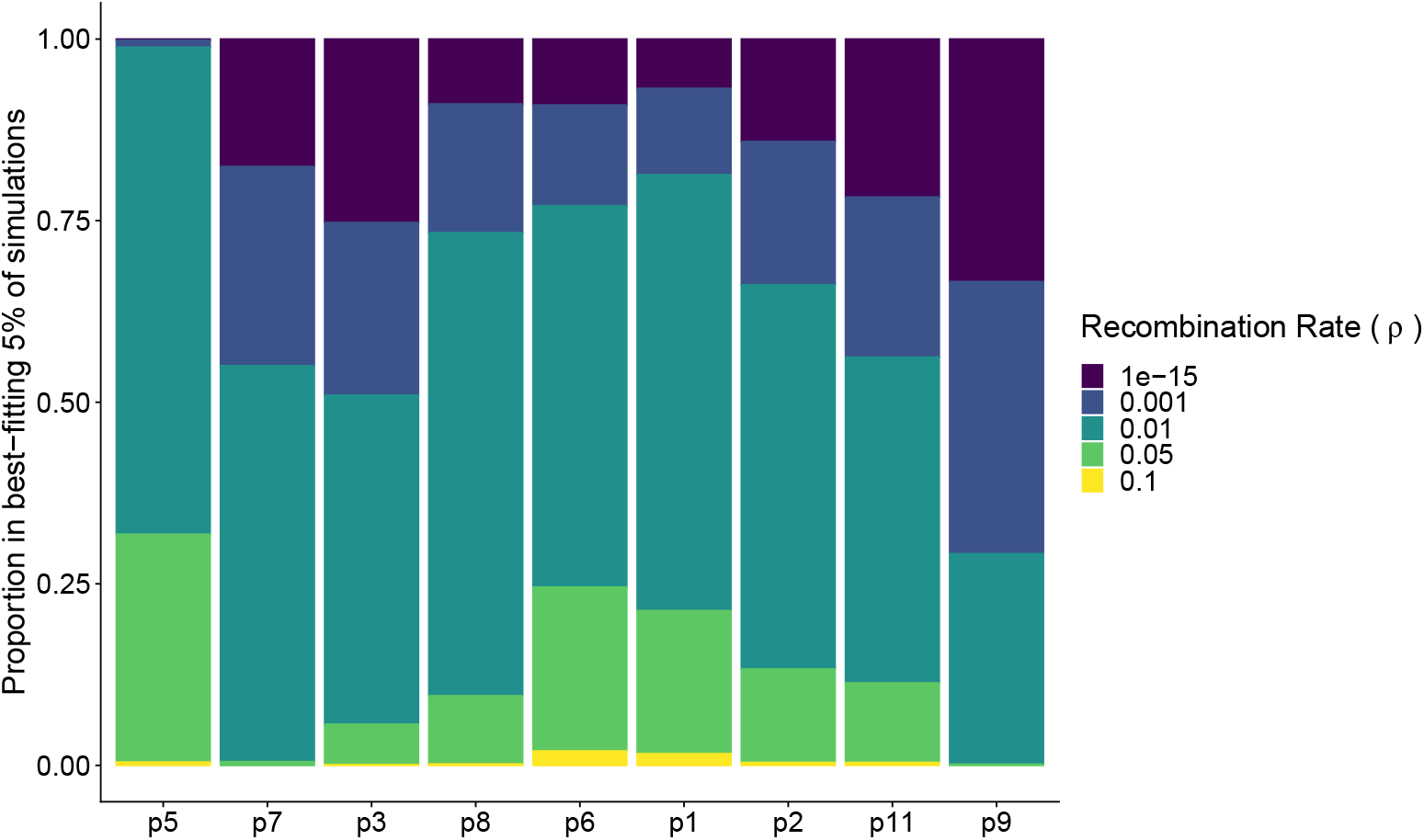
The recombination breakdown within the best-fitting 5% of simulations for each patient. For this figure, we bootstrapped the scores within each recombination strata so that each strata contained 10,000 values. For each bootstrap replicate, we normalized the score across the 50,000 samples, and calculated the relative frequency of each recombination rate in the top 5% of matching simulations. Here, we plot the mean relative frequency of each recombination strata over 100 bootstrap replicates. Patients are organized by the mean of the normalized top scores, with patient 5 having the lowest (best-fitting) mean score.

**Figure 8:**
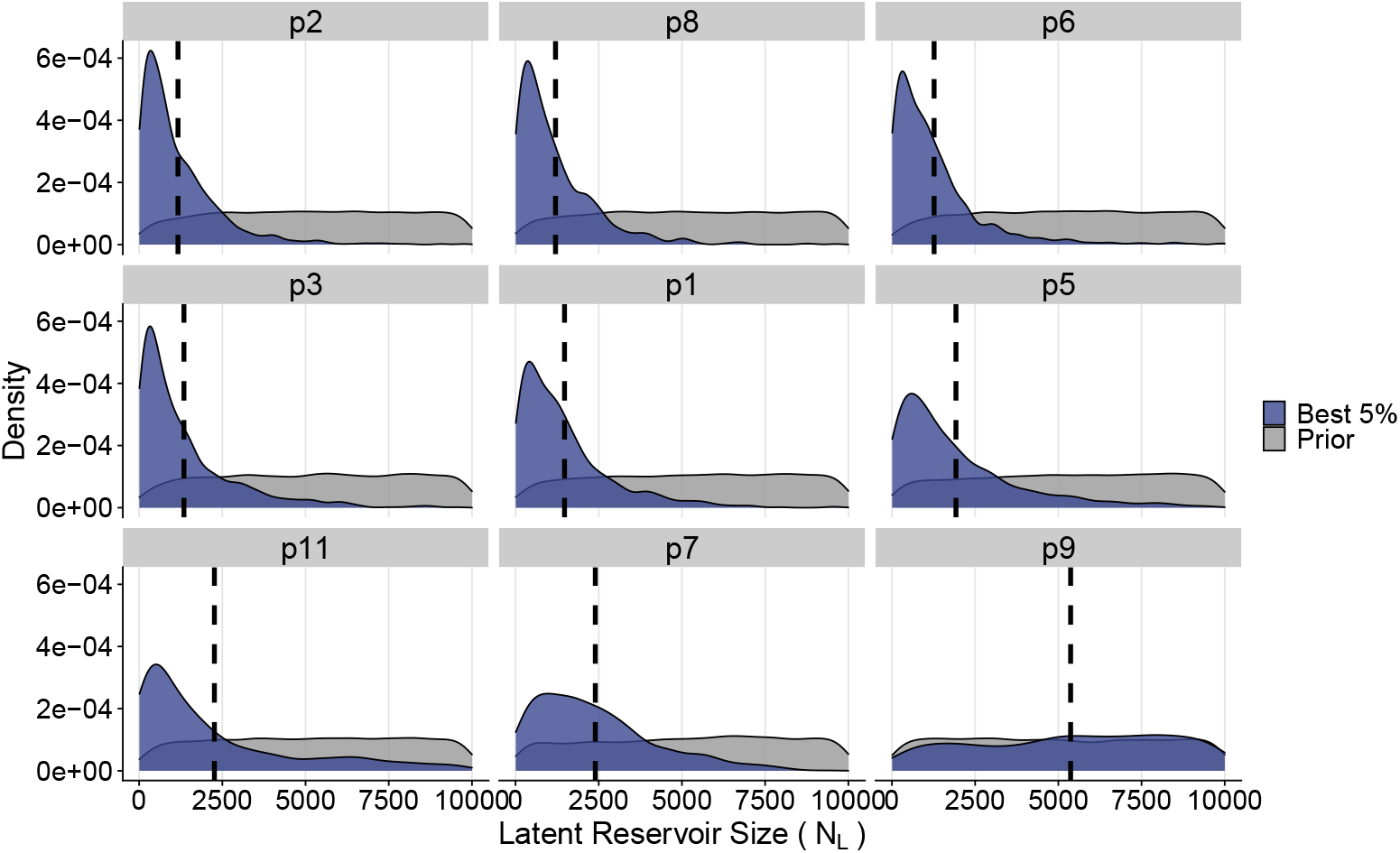
Marginal distributions of *N*_*L*_ across the nine patients. Colors distinguish the density of parameters between simulations with the best-fitting 5% of distance scores (blue) and the highest 95% of distance scores (grey). The dashed line indicates the mean of the best 5% distribution. Patients are ordered by increasing mean latent reservoir size of the best-fitting 5% of simulations. This figure represents the results of one of the bootstrap replicates.

## Discussion

The importance of recombination and the latent reservoir in within-host HIV-1 evolutionary dynamics has been described in detail since the mid 1990’s [49, 50]. Although these two processes have been modeled individually and together using forward mathematical models and simulations [47, 51–53], both processes have been largely ignored in phylodynamic investigations because of the theoretical and computational problems in jointly inferring the phylogeny and recombination patterns. As efforts continue towards finding a functional cure of HIV-1 that involves eliminating or reducing the size of the latent reservoir, studying the interaction between the latent reservoir and recombination remains key. In this study we extended a previously developed within-host coalescent model [12] into an ARG simulation model that allows for lineages to coalesce, recombine, and cycle in and out of a latent state. We found that these complex evolutionary dynamics leave tractable signals in bifurcating trees reconstructed using hierarchical clustering which can be quantified in readily measurable topological and distance features. Additionally, we used a sampling estimation framework to evaluate the intensity of recombination and the latent reservoir size in HIV-1 patient trees.

Our within-host HIV-1 ARG simulator produced trees that qualitatively and quantitatively captured HIV-1 within-host evolutionary dynamics. The topological and distance-based statistics from reconstructed trees were within the range of empirical HIV-1 phylogenies, and our distance matching algorithm showed the ability to recover distinguishing tree characteristics when applied to patients with visually distinct evolutionary histories. In general, there was a positive correlation between the intensity of recombination and the size of the latent reservoir. Here, our model assumes that the half-life of the latent reservoir is fixed such that large latent pools will produce greater number of activation events, and consequently, faster recombination rates are required to maintain a ladder-like tree shape. Because our model lacks a fitness cost for newly activated latent viruses, however, it is possible that we are overstating this correlation as re-activated latent cells that do not recombine with contemporary virus are not likely to survive long enough to be sampled at high frequency in reality [47]. Nevertheless, we can derive conclusions and generate hypotheses about the relative strength and consistency of the recombination rate and size of the latent reservoir. We found consistency between the recombination rates that produce the best-fitting trees to the empirical data and the current biological estimates of the effective recombination rate. The estimated effective recombination rate, 1.4 × 10^−5^ to 1.38 × 10^−4^ per base per generation, corresponds to a rate of 0.008 − 0.081 recombination events per lineage per day in our simulation of a 700 nt genomic fragment.

On the latent reservoir size, we found support for latent reservoirs that are on the order of 10^2^-103. Large latent reservoirs closer to 10^4^ produce trees with extreme external to internal branch ratios (Figure 4), at times 100x larger than the maximum empirical measurements (Figure S3). A 10^2^ latent reservoir corresponds to the estimate of 1 per 10^6^ million CD4+ T cells. While our results suggest the latent reservoir is near the smaller end of the estimated spectrum and might vary between individuals, there remain unanswered questions relating the absolute size of the reservoir with how much virus it produces and quantifying the adaptive impacts it has on generating genetic diversity available for immune escape.

The third biological feature we included in our within-host ARG simulation model was the population demography in the form of (internal) bottlenecks. The lack of individual-level fitness differences between lineages might explain the tendency of our model to produce somewhat more balanced trees than what may be seen in real HIV-1 trees. In our model, the coalescence rates of lineages are neutral between bottlenecks. In reality, low-level positive selection from the immune system would create fitness differences between lineages, particularly for reactivated latent lineages that would be less fit because of long-term immunological memory [54–56]. These fitness differences would likely increase the ladder-like structure of a tree. In addition to creating more balanced trees, this neutrality likely causes the variability in possible evolutionary histories for a given Θ, particularly when recombination is low. Lastly, this model simplification might be creating tension in the distance scoring. Because the Sackin index of simulated trees was often far below the empirical value, when the Sackin index is close, it has a disproportionate weight on the overall scoring. To address these shortcomings, future work should consider alternative formulations of changes in demography.

In our distance algorithm we introduced the ranked distance matrix as a new statistical probe, which has two notable features. First, because the ARG models the time between events, the distance matrix obtained from an ARG-decomposed bifurcating tree accounts for the true evolutionary distance in time between the extant tips. Second, distance matrices can be computed from sequence data directly without first inferring a phylogenetic tree that may impose artifactual structures on the data in the context of high levels of recombination. However, a potential source of error comes from matching the time scale of the ARG to the genetic distance of the sequence data. In our formulation, an optimized scaling factor transforms the time distance to match the genetic distance and is independent of any specific genome-region. However, the scaling factor assumes a constant rate over the course of an individual’s infection and assumes that multiple mutations at an individual site have been perfectly accounted for.

In conclusion, we have developed a novel HIV-1 within-host simulator that includes realistic features such as recombination, latency, and internal bottlenecks. These processes generate an ARG, a complex evolutionary structure formed by loops, mixed time-signals, and unbalanced growth. Because such structures are, at this time, exceedingly difficult to infer from real data, we also developed an ARG decomposition method to visualize this complex structure as a familiar bifurcating tree. Together, these tools makes it possible to match recombination, latency, and bottleneck effects to real HIV-1 within-host data, where the combined effects otherwise could not be analyzed.

## supplemental

Supplement: Recombination smooths the time-signal disrupted by latency in within-host HIV phylogenies

## Acknowledgements

This study was supported by the NIH/NIAID grant R01AI087520 to TL. LAC was supported by the Department of defense (DoD) through the National Defense Science & Engineering Graduate Fellowship (NDSEG) Program. Los Alamos National Laboratory is operated by Triad National Security, LLC, for the National Nuclear Security Administration of U.S. Department of Energy (Contract No. 89233218CNA000001). The content is solely the responsibility of the authors and does not represent the official views of the sponsors.

## Code Availability

The coalescent ARG simulator and ARG decomposition method are available at https://github.com/MolEvolEpid/ARG_simul-decomp.

---

**Figure S1:**
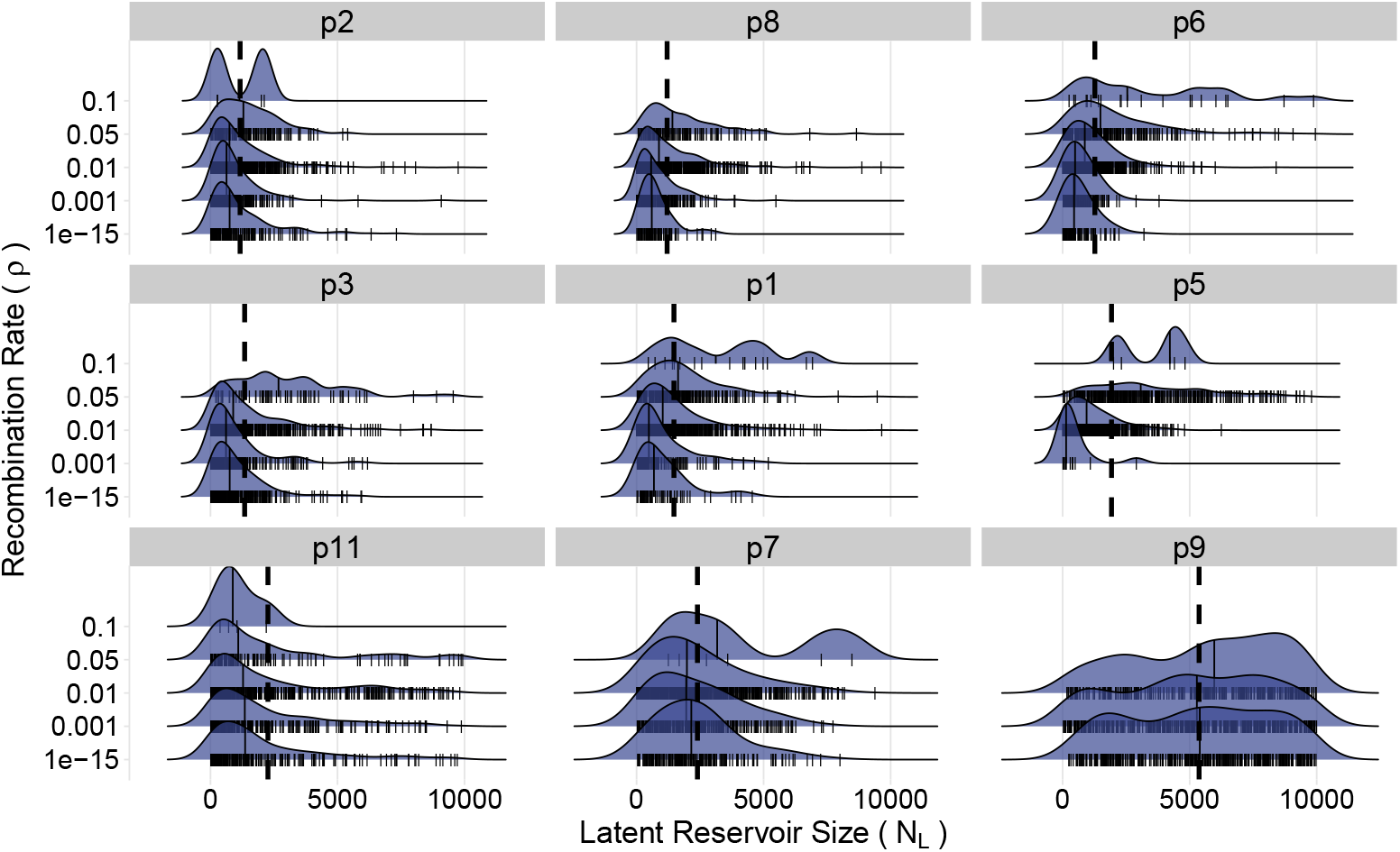
Distribution of latent pool size stratified by recombination rate. For each patient, the posterior samples for reservoir size are shown stratified by recombination rate. The dashed line shows the maximum posterior estimated of the reservoir size for each patient. Patients are ordered by increasing mean latent reservoir size of the best-fitting 5% of simulations.

**Figure S2:**
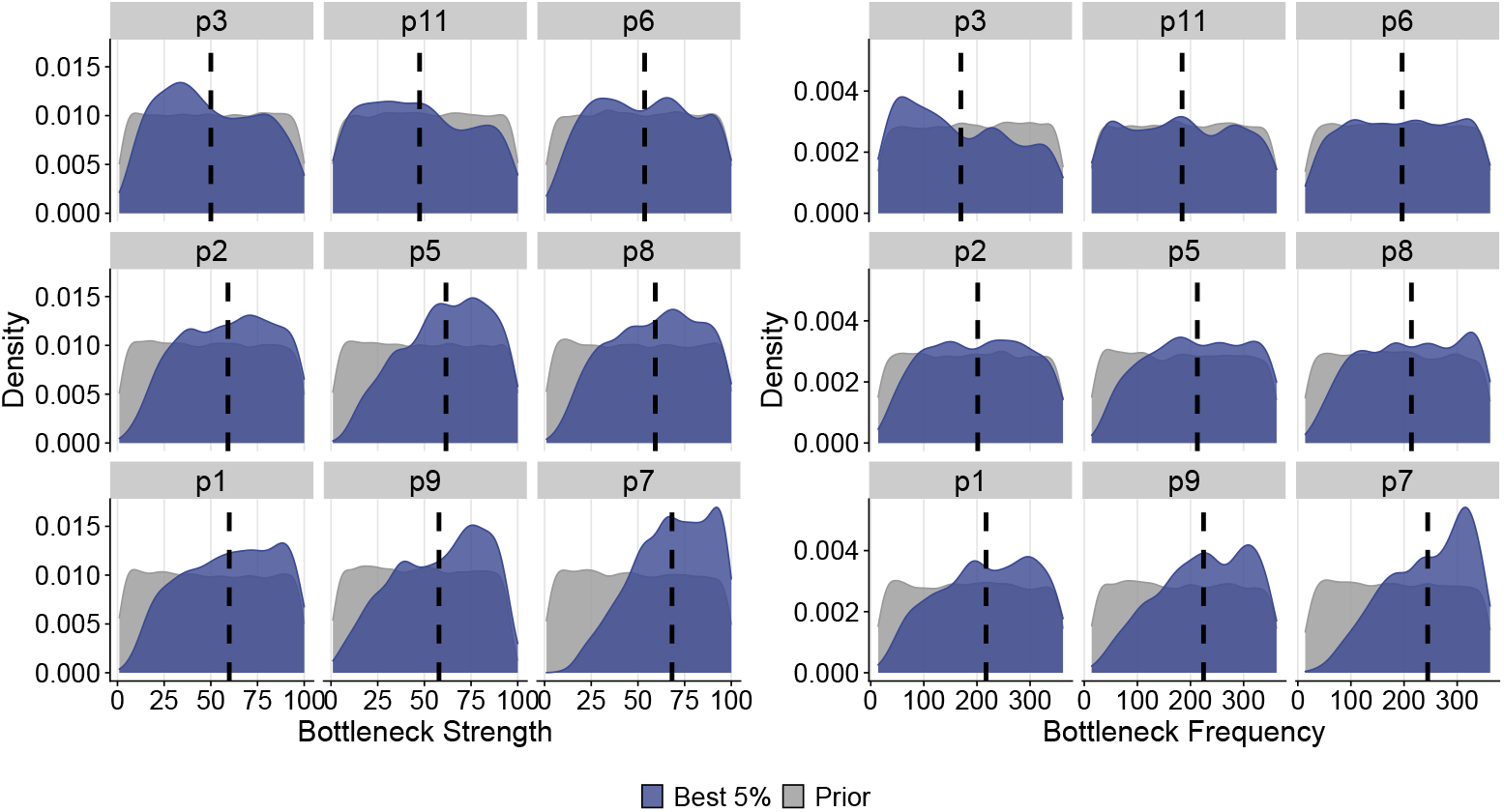
Marginal distributions of the bottleneck attributes of strength and frequency across the nine patients. Colors distinguish the density of parameters between simulations with the best (lowest) 5% of distance scores (red) and the highest 95% of distance scores (blue). The dashed line indicates the mean of the best-fitting 5% of simulations. Patients are ordered by the best-fitting 5% mean of bottleneck frequency values.

**Figure S3:**
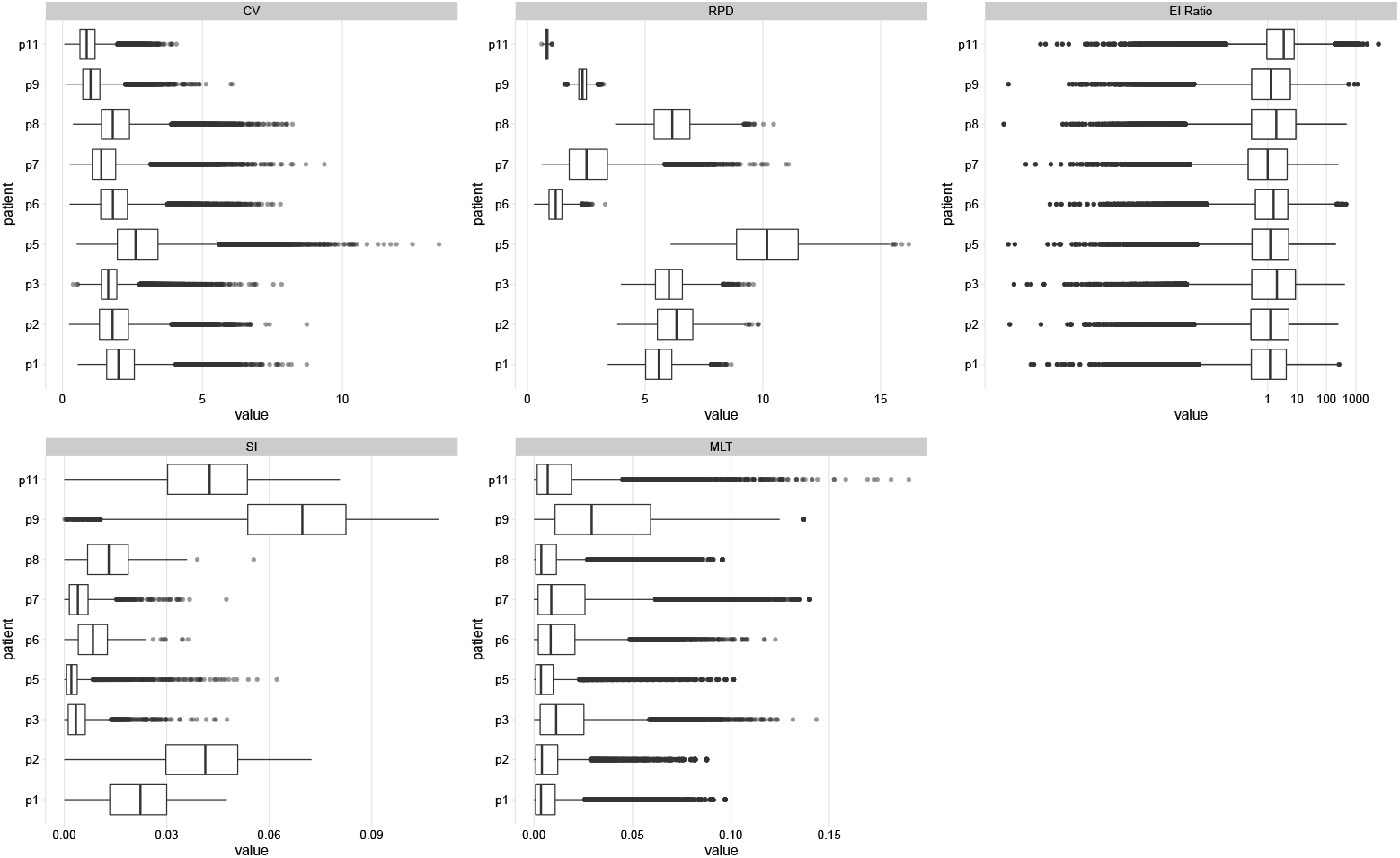
Distributions of topological and distance statistics derived from the complete collection of the ARG-simulated reconstructed phylogenies for each patient.

## Notes

### Competing Interest Statement

The authors have declared no competing interest.

## Literature Cited

[1] Volz, E. M., E. Romero-Severson, T. Leitner. 2017. Phylodynamic Inference across Epidemic Scales. Molecular Biology and Evolution 34:1276–1288.

[2] Leitner, T. 2019. Phylogenetics in HIV transmission: taking within-host diversity into account. Current opinion in HIV and AIDS 14:181–187.

[3] Mendes, F. K., A. P. Livera, M. W. Hahn. 2019. The perils of intralocus recombination for inferences of molecular convergence. Philosophical Transactions of the Royal Society B 374:20180244.

[4] Immonen, T. T., T. Leitner. 2014. Reduced evolutionary rates in HIV-1 reveal extensive latency periods among replicating lineages. Retrovirology 11:81.

[5] Kouyos, R., V. von Wyl, S. Yerly, et al. 2010. Molecular Epidemiology Reveals Long-Term Changes in HIV Type 1 Subtype B Transmission in Switzerland. The Journal of Infectious Diseases 201:1488– 1497.

[6] Volz, E. M., E. Ionides, E. O. Romero-Severson, et al. 2013. HIV-1 Transmission during Early Infection in Men Who Have Sex with Men: A Phylodynamic Analysis. PLoS medicine 10:1–12.

[7] Grabowski, M. K., J. Lessler, A. D. Redd, et al. 2014. The Role of Viral Introductions in Sustaining Community-Based HIV Epidemics in Rural Uganda: Evidence from Spatial Clustering, Phylogenetics, and Egocentric Transmission Models. PLoS medicine 11.

[8] Fisher, M., D. Pao, A. E. Brown, et al. 2010. Determinants of HIV-1 transmission in men who have sex with men: a combined clinical, epidemiological and phylogenetic approach. AIDS 24:1739–1747.

[9] Stadler, T., D. Kühnert, S. Bonhoeffer, et al. 2013. Birth-death skyline plot reveals temporal changes of epidemic spread in HIV and hepatitis C virus (HCV). Proceedings of the National Academy of Sciences of the United States of America 110:228–33.

[10] Rasmussen, D. A., E. M. Volz, K. Koelle. 2014. Phylodynamic inference for structured epidemiological models. PLoS computational biology 10:e1003570.

[11] Ratmann, O., A. Van Sighem, D. Bezemer, et al. 2016. Sources of HIV infection among men having sex with men and implications for prevention. Science translational medicine 8.

[12] Romero-Severson, E., H. Skar, I. Bulla, et al. 2014. Timing and Order of Transmission Events Is Not Directly Reflected in a Pathogen Phylogeny. Molecular Biology and Evolution 39:2472–2482.

[13] Giardina, F., E. O. Romero-Severson, J. Albert, et al. 2017. Inference of Transmission Network Structure from HIV Phylogenetic Trees. PLoS computational biology 13:e1005316.

[14] Hall, M. D., C. Colijn. 2019. Transmission Trees on a Known Pathogen Phylogeny: Enumeration and Sampling. Molecular Biology and Evolution 36:1333–1343.

[15] Siepel, A. C., A. L. Halpern, C. Macken, et al. 1995. A computer program designed to screen rapidly for hiv type 1 intersubtype recombinant sequences. AIDS research and human retroviruses 11:1413– 1416.

[16] Salminen, M. O., J. K. Carr, D. S. Burke, et al. 1995. Identification of breakpoints in intergenotypic recombinants of HIV type 1 by bootscanning. AIDS research and human retroviruses 11:1423–1425.

[17] Giorgi, E. E., B. Funkhouser, G. Athreya, et al. 2010. Estimating time since infection in early homogeneous hiv-1 samples using a poisson model. BMC bioinformatics 11:1–7.

[18] Leitner, T., E. Romero-Severson. 2018. Phylogenetic patterns recover known HIV epidemiological relationships and reveal common transmission of multiple variants. Nature microbiology 3:983–988.

[19] Song, H., E. E. Giorgi, V. V. Ganusov, et al. 2018. Tracking HIV-1 recombination to resolve its contribution to hiv-1 evolution in natural infection. Nature communications 9:1–15.

[20] Neher, R. A., T. Leitner. 2010. Recombination Rate and Selection Strength in HIV Intra-patient Evolution. PLoS computational biology 6:e1000660.

[21] Batorsky, R., M. F. Kearney, S. E. Palmer, et al. 2011. Estimate of effective recombination rate and average selection coefficient for HIV in chronic infection. Proceedings of the National Academy of Sciences of the United States of America 108:5661–6.

[22] Josefsson, L., S. Palmer, N. R. Faria, et al. 2013. Single Cell Analysis of Lymph Node Tissue from HIV-1 Infected Patients Reveals that the Majority of CD4+ T-cells Contain One HIV-1 DNA Molecule. PLoS pathogens 9:e1003432.

[23] Mansky, L. M., H. M. Temin. 1995. Lower in vivo mutation rate of human immunodeficiency virus type 1 than that predicted from the fidelity of purified reverse transcriptase. Journal of Virology 69:5087–94.

[24] Gao, F., Y. Chen, D. N. Levy, et al. 2004. Unselected mutations in the human immunodeficiency virus type 1 genome are mostly nonsynonymous and often deleterious. Journal of Virology 78:2426–33.

[25] Fisher, R. 1930. The general theory of natural selection. Oxford: Clarendon.

[26] Muller, H. J. 1932. Some Genetic Aspects of Sex. The American naturalist 66:118–138.

[27] Crow, J. F., M. Kimura. 1965. Evolution in sexual and asexual population. The American naturalist 99:439–450.

[28] Felsenstein, J. 1974. The evolutionary advantage of recombination. Genetics 78.

[29] Michod, R. E., H. Bernstein, A. M. Nedelcu. 2008. Adaptive value of sex in microbial pathogens. Journal of molecular epidemiology and evolutionary genetics in infectious diseases 8:267–285.

[30] Kellam, P., B. A. Larder. 1995. Retroviral recombination can lead to linkage of reverse transcriptase mutations that confer increased zidovudine resistance. Journal of Virology 69:669–74.

[31] Moutouh, L., J. Corbeil, D. D. Richman. 1996. Recombination leads to the rapid emergence of HIV1 dually resistant mutants under selective drug pressure. Proceedings of the National Academy of Sciences of the United States of America 93:6106–11.

[32] Gu, Z., Q. Gao, E. A. Faust, et al. 1995. Possible involvement of cell fusion and viral recombination in generation of human immunodeficiency virus variants that display dual resistance to AZT and 3TC. The Journal of general virology 76:2601–2605.

[33] Kondrashov, F. A., A. S. Kondrashov. 2001. Multidimensional epistasis and the disadvantage of sex. Proceedings of the National Academy of Sciences of the United States of America 98:12089–12092.

[34] Moradigaravand, D., R. Kouyos, T. Hinkley, et al. 2014. Recombination Accelerates Adaptation on a Large-Scale Empirical Fitness Landscape in HIV-1. PLoS genetics 10:e1004439.

[35] Griffiths, R., P. Marjoram. 1996. Ancestral Inference from Samples of DNA Sequences with Recombination. Journal of computational biology 3:479–502.

[36] Hein, J. 1990. Reconstructing evolution of sequences subject to recombination using parsimony. Mathematical Biosciences 98:185–200.

[37] Rasmussen, M. D., M. J. Hubisz, I. Gronau, et al. 2014. Genome-Wide Inference of Ancestral Recombination Graphs. PLOS genetics 10:e1004342.

[38] Wang, L., K. Zhang, L. Zhang. 2001. Perfect phylogenetic networks with recombination. Journal of computational biology 8:69–78.

[39] Romero-Severson, E., G. Meadors, E. Volz. 2014. A Generating Function Approach to HIV Transmission with Dynamic Contact Rates. Mathematical Modelling of Natural Phenomena 9:121–135.

[40] Wakeley, J. 2009. Coalescent theory: an introduction. Roberts & Co. Publishers, Greenwood Village, Colo.

[41] Siliciano, J. D., J. Kajdas, D. Finzi, et al. 2003. Long-term follow-up studies confirm the stability of the latent reservoir for HIV-1 in resting CD4+T cells. Nature Med. 9:727–728.

[42] Csardi, G. 2019. Package ‘igraph’ Title Network Analysis and Visualization.

[43] Desper, R., O. Gascuel. 2002. Fast and Accurate Phylogeny Reconstruction Algorithms Based on the Minimum-Evolution Principle. 357–374. Springer, Berlin, Heidelberg.

[44] Shankarappa, R., J. B. Margolick, S. J. Gange, et al. 1999. Consistent Viral Evolutionary Changes Associated with the Progression of Human Immunodeficiency Virus Type 1 Infection Downloaded from. Tech. Rep. 12.

[45] Tamura, K., M. Nei. 1993. Estimation of the number of nucleotide substitutions in the control region of mitochondrial DNA in humans and chimpanzees. Molecular biology and evolution 10:512–526.

[46] Zanini, F., J. Brodin, L. Thebo, et al. 2015. Population genomics of intrapatient HIV-1 evolution. Elife 4.

[47] Immonen, T. T., J. M. Conway, E. O. Romero-Severson, et al. 2015. Recombination Enhances HIV-1 Envelope Diversity by Facilitating the Survival of Latent Genomic Fragments in the Plasma Virus Population. PLoS computational biology.

[48] Skar, H., R. N. Gutenkunst, K. Wilbe Ramsay, et al. 2011. Daily sampling of an hiv-1 patient with slowly progressing disease displays persistence of multiple env subpopulations consistent with neutrality. PLoS One 6:e21747.

[49] Finzi, D., M. Hermankova, T. Pierson, et al. 1997. Identification of a Reservoir for HIV-1 in Patients on Highly Active Antiretroviral Therapy. Science (80-.). 278:1295–1300.

[50] Ruelas, D. S., W. C. Greene. 2013. An Integrated Overview of HIV-1 Latency. Cell 155:519–529.

[51] Hill, A. L. 2017. Mathematical Models of HIV Latency. Current topics in microbiology and immunology.

[52] Murray, J. M., J. Zaunders, S. Emery, et al. 2017. HIV dynamics linked to memory CD4+ T cell homeostasis. PLoS One.

[53] Doekes, H. M., C. Fraser, K. A. Lythgoe. 2017. Effect of the Latent Reservoir on the Evolution of HIV at the Within- and Between-Host Levels. PLoS computational biology.

[54] Wei, X., J. M. Decker, S. Wang, et al. 2003. Antibody neutralization and escape by HIV-1. Nature 422:307–312.

[55] Richman, D. D., T. Wrin, S. J. Little, et al. 2003. Rapid evolution of the neutralizing antibody response to HIV type 1 infection. Proceedings of the National Academy of Sciences of the United States of America 100:4144–9.

[56] Bunnik, E. M., L. Pisas, A. C. van Nuenen, et al. 2008. Autologous neutralizing humoral immunity and evolution of the viral envelope in the course of subtype B human immunodeficiency virus type 1 infection. Journal of Virology 82:7932–41.

[57] Jetzt, A. E., H. Yu, G. J. Klarmann, et al. 2000. High Rate of Recombination throughout the Human Immunodeficiency Virus Type 1 Genome. Journal of Virology 74:1234–1240.

[58] Zhuang, J., A. E. Jetzt, G. Sun, et al. 2002. Human immunodeficiency virus type 1 recombination: rate, fidelity, and putative hot spots. Journal of Virology 76:11273–82.

[59] Chun, T. W., L. Stuyver, S. B. Mizell, et al. 1997. Presence of an inducible HIV-1 latent reservoir during highly active antiretroviral therapy. Proceedings of the National Academy of Sciences of the United States of America 94:13193–7.

[60] Wong, J. K., M. Hezareh, H. F. Günthard, et al. 1997. Recovery of Replication-Competent HIV Despite Prolonged Suppression of Plasma Viremia. Science (80-.). 278:1291–1295.

[61] Ho, Y.-C., L. Shan, N. Hosmane, et al. 2013. Replication-Competent Noninduced Proviruses in the Latent Reservoir Increase Barrier to HIV-1 Cure. Cell 155:540–551.

[62] Janzen, T., S. Höhna, R. S. Etienne. 2015. Approximate Bayesian Computation of diversification rates from molecular phylogenies: introducing a new efficient summary statistic, the nLTT. Methods in ecology and evolution 6:566–575.

